# Chronic alcohol consumption alters sex-dependent BNST neuron function in rhesus macaques

**DOI:** 10.1101/2024.04.11.589120

**Authors:** Kristen E. Pleil, Kathleen A. Grant, Verginia C. Cuzon Carlson, Thomas L. Kash

## Abstract

Repeated alcohol drinking contributes to a number of neuropsychiatric diseases, including alcohol use disorder and co-expressed anxiety and mood disorders. Women are more susceptible to the development and expression of these diseases with the same history of alcohol exposure as men, suggesting they may be more sensitive to alcohol-induced plasticity in limbic brain regions controlling alcohol drinking, stress responsivity, and reward processing, among other behaviors. Using a translational model of alcohol drinking in rhesus monkeys, we examined sex differences in the basal function and plasticity of neurons in the bed nucleus of the stria terminalis (BNST), a brain region in the extended amygdala shown to be a hub circuit node dysregulated in individuals with anxiety and alcohol use disorder. We performed slice electrophysiology recordings from BNST neurons in male and female monkeys following daily “open access” (22 hr/day) to 4% ethanol and water for more than one year or control conditions. We found that BNST neurons from control females had reduced overall current density, hyperpolarization-activated depolarizing current (I_h_), and inward rectification, as well as higher membrane resistance and greater synaptic glutamatergic release and excitatory drive, than those from control males, suggesting that female BNST neurons are more basally excited than those from males. Chronic alcohol drinking produced a shift in these measures in both sexes, decreasing current density, I_h_, and inward rectification and increasing synaptic excitation. In addition, network activity-dependent synaptic inhibition was basally higher in BNST neurons of males than females, and alcohol exposure increased this in both sexes, a putative homeostatic mechanism to counter hyperexcitability. Altogether, these results suggest that the rhesus BNST is more basally excited in females than males and chronic alcohol drinking produces an overall increase in excitability and synaptic excitation. These results shed light on the mechanisms contributing to the female-biased susceptibility to neuropsychiatric diseases including co-expressed anxiety and alcohol use disorder.

## 1. Introduction

Excessive alcohol consumption is a leading risk factor for many negative health outcomes, including neuropsychiatric diseases such as alcohol use disorder (AUD) and anxiety disorders that are prevalent and commonly co-expressed (Kessler et al., 2005; Smith and Randall, 2012; Smyth et al., 2015; Gimeno et al., 2017). Chronic alcohol exposure leads to a myriad of adaptations in critical circuit nodes that regulate various functions including stress reactivity, reward sensitivity, affect, and cognition (Koob and Volkow, 2016; Volkow et al., 2016; Uhl et al., 2019; Guinle and Sinha, 2020; Goldfarb et al., 2022; Sinha, 2022). The bed nucleus of the stria terminalis (BNST) is a component of the extended amygdala that is densely interconnected with hindbrain and midbrain regions that receives input about the environment and internal state of the individual and cortical regions important for behavioral control (Dong et al., 2001; Dong and Swanson, 2004; Dong et al., 2017; Vranjkovic et al., 2017). As such, human neuroimaging studies have shown that the BNST is a hub region in the dysregulated neural circuitry found in both AUD and anxiety (O’Daly et al., 2012; Avery et al., 2014; Avery et al., 2016) that may play a causal role in the expression of alcohol drinking and anxiety behaviors. Work from our group and others has found that the BNST is a critical site for the regulation of excessive alcohol consumption in rodents (Eiler et al., 2003; Eiler and June, 2007; Pina et al., 2015; Pleil et al., 2015b; Rinker et al., 2016); and, many studies have described plasticity in the excitability and synaptic transmission of BNST neurons following chronic alcohol exposure related to the maintenance of excessive alcohol consumption and withdrawal-induced anxiety and negative affect (Olive et al., 2002; Kash et al., 2009; McElligott and Winder, 2009; Kash, 2012; McElligott et al., 2013; Silberman et al., 2013; Kash et al., 2015; Pleil et al., 2015a).

Women are more sensitive to the negative health outcomes associated with acute and chronic alcohol intake and are more susceptible to co-development and co-expression of these neuropsychiatric diseases than men (Grant et al., 2004a; Grant et al., 2004b; Kessler et al., 2005; Peltier et al., 2019; Flores-Bonilla and Richardson, 2020; Guinle and Sinha, 2020). In addition, females transition from social to abusive alcohol drinking related to AUD more rapidly than males, and heavy alcohol drinking is increasing at an alarming pace in women, especially since the beginning of the Covid-19 pandemic (Cheng et al., 2016; Pollard et al., 2020). Together, these observations suggest that females may be more sensitive to alcohol-induced plasticity in the BNST that promotes alcohol drinking and anxiety. We recently found in mice that stress-sensitive subpopulations of BNST neurons that regulate alcohol drinking and anxiety behavior display basal sex differences in their function, including higher basal excitability and excitatory synaptic transmission in females than males. Further, we found that repeated alcohol drinking increased these measures in males to female-like levels (Levine et al., 2021). In the current study, we evaluate sex-dependent plasticity in BNST neurons using a rhesus monkey model of chronic alcohol self-administration that captures the spectrum of human alcohol consumption, including individual variability in early patterns of drinking that predicts long-term alcohol intakes (Grant et al., 2008; Baker et al., 2014; Baker et al., 2017). In this model, males with a history of chronic voluntary alcohol consumption show altered function of BNST neurons through adaptations in inhibitory synaptic transmission and passive membrane properties (Pleil et al., 2016). Here, we expand upon those findings by examining sex differences in electrophysiological properties, voltage-current relationships, and network activity dependent/independent excitatory and inhibitory synaptic transmission in the rhesus BNST following chronic alcohol consumption. These comparative experiments identify critical sex differences in and effects of alcohol on BNST neuron function that have implications for sexually dimorphic outcomes from chronic alcohol drinking in humans.

## 2. Materials and Methods

### 2.1 Subjects

Male and female rhesus monkeys (*Macaca mulatta*) bred and raised at the Oregon National Primate Research Center of the Oregon Health & Science University underwent a well-documented protocol of alcohol induction and chronic alcohol self-administration (**Fig. 1A**) or control procedure as previously described (Grant et al., 2008; Baker et al., 2014; Baker et al., 2017). Detailed drinking and other behavioral measures from these cohorts’ subjects (cohorts female 6b, male 7a, and male 7b) are available by request from the Monkey Alcohol Tissue Research Resource (MATRR.com). Briefly, monkeys within a cohort were singly housed, with partner-pairing 1-2 hrs/day, in a colony room and trained to use an operant panel to deliver food and water. They then were trained to self-administer water (at a volume equivalent to 1.5g/kg ethanol (EtOH) and then escalating doses of 4% EtOH (w/v) (0.5 g/kg, 1.0 g/kg, then 1.5 g/kg daily) in 30-day epochs, followed by a 12-month period of open access (22 hr/day) to 4% EtOH in which alcohol-drinking males (EtOH M, N = 10) and females (EtOH F, N = 5) had similar average daily intake (2.37 g/kg and 2.16 g/kg, respectively; **Fig. 1B**). A control group of males (CON M, N = 7) and a control group of females (CON F, N = 3) that did not have EtOH access, but was provided calorically matched maltose, were used for comparison.

**Figure 1:**
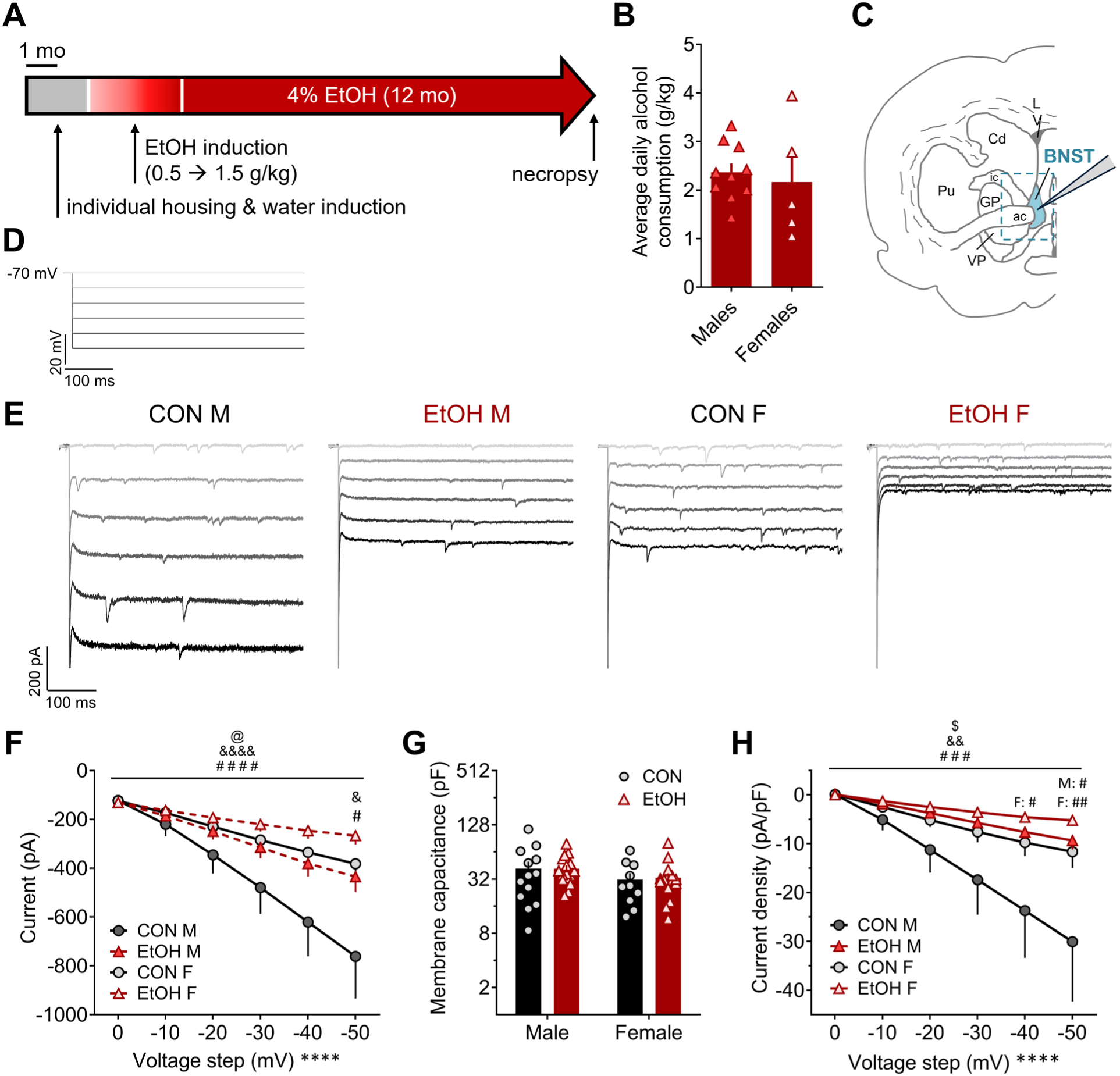
Sex-dependent current responses to membrane hyperpolarization in monkey BNST neurons during voltage clamp recordings is modulated by a history of chronic voluntary alcohol intake. A) Voluntary alcohol access paradigm in male and female rhesus monkeys, in which animals were singly housed and introduced to interactive panels to deliver water (∼6 months) and then alcohol to achieve escalating intake up to 1.5 g/kg (three months), followed by open access to alcohol for 12 months prior to necropsy and electrophysiological recordings. **B)** Average daily 4% EtOH intake across the open access period in monkeys used in this study was similar between sexes (t_13_ = 0.45, p > 0.65). **C)** Schematic of a monkey brain on the coronal plane containing BNST, with blue dashed outline representing the approximate dissected tissue used for slice electrophysiology recordings. **D)** Stimulus waveform for electrophysiological recordings measuring the current-voltage relationship in BNST neurons from control males (CON M), alcohol males (EtOH M), control females (CON F), and alcohol females (EtOH F), in which cells held under voltage clamp at a holding potential of -70 mV were underwent increasingly hyperpolarizing steps of 10 mV to -120 mV. E) Representative traces for current responses to the stimulus in D. **F)** Both sex and chronic alcohol drinking altered the raw current response to hyperpolarization of the membrane potential. 3xRM-ANOVA: main effects of sex (F_1,51_ = 5.15, ^@^p = 0.028) and voltage step (F_5,255_ = 65.78, ****p < 0.0001) and a trend of EtOH (F_1,51_ = 3.10, p = 0.085); interactions between sex x step (F_5,255_ = 11.30, ^&&&&^p < 0.0001) and EtOH x step (F_5,255_ = 7.70, ^####^p < 0.0001) but no others (ps > 0.10). Multiple t-tests with H-S corrections across all 6 voltage steps showed that CON M and CON F differed on the -50 mV step (t_126_ = 2.95, ^&^p = 0.023) and CON M and EtOH M differed on the -50 mV step (t_162_ = 3.03, ^#^p = 0.017); all other comparisons, including between CON F and EtOH F, did not differ (ps > 0.10). G) Membrane capacitance was unaffected by sex or EtOH. 2xANOVA:no effects of sex, EtOH, or their interaction (ps > 0.10). **H)** Current density responses to membrane hyperpolarization (change in current normalized to cell capacitance) were affected by sex and EtOH. 3xRM-ANOVA: main effects of EtOH (F_1,51_ = 4.77, ^$^p = 0.034) and voltage step (F_5,255_ = 21.11, ****p < 0.0001) and a trend of sex (F_1,51_ = 2.94, p = 0.093); interactions between sex x step (F_5,255_ = 3.39, ^&&^p = 0.006) and EtOH x step (F_5,255_ = 4.86, ^###^p =0.0003) but no others (ps > 0.25). Multiple t-tests with H-S corrections across all six voltage steps showed that CON M and EtOH M differed on the -50 mV step (t_162_ = 3.10, ^#^p = 0.013) with a trend on the -40 mV step (t_162_ = 2.41, p = 0.083); CON F and EtOH F differed on the -50 mV step (t_144_ = 3.41, ^##^p = 0.005) and -40 mV step (t_144_ = 2.76, ^#^p = 0.033); all other comparisons, including between CON M and CON F, did not differ (ps > 0.10).

### 2.2 Necropsy and slice preparation

Monkeys underwent necropsy under ketamine (10 mg/kg) plus isoflurane at a dose to maintain a deep anesthetic plane in the morning at the time they usually were given alcohol access (Daunais et al., 2010). Transcardial perfusion was performed with ice-cold oxygenated artificial cerebrospinal fluid (aCSF), and a craniotomy was conducted. A 4 x 6 x 8 mm unilateral coronal block of fresh brain tissue containing the BNST was collected (as outlined in a dashed blue line in **Fig. 1b**), transferred to ice-cold oxygenated (95% O_2_/5% CO_2_) aCSF containing (in mM) 124 NaCl, 4.5 KCl, 1 MgCl_2_, 26 NaHCO_3_, 1.2 NaH_2_PO_4_, 10 glucose and 2 CaCl_2_, and transported on ice for slicing. The tissue block was transferred to ice-cold oxygenated sucrose aCSF (saCSF) containing (in mM) 194 sucrose, 30 NaCl, 4.5 KCl, 1 MgCl_2_, 26 NaHCO_3_, 1.2 NaH_2_PO4 and 10 glucose, and it was sectioned into 250 μm coronal slices on a VT 1200S vibratome (Leica Biosystems, Wetzlar, Germany). Slices were transferred to aCSF in a holding chamber held at 37°C for one hr and then subsequently maintained at room temperature until used for slice electrophysiology.

### 2.3 Slice electrophysiology

Whole-cell patch-clamp slice electrophysiological recordings were performed as previously described (Pleil et al., 2012; Pleil et al., 2015a; Pleil et al., 2015b; Pleil et al., 2016). Individual slices were transferred to a recording chamber within the fixed stage of an upright microscope, secured with platinum weights, and constantly perfused with ∼30°C aCSF. BNST neurons were identified under differential interference contrast (DIC) using an infrared camera with 40x water-immersion objective and digital computer image. Borosilicate glass capillaries (1.5 mm outer diameter, 0.86 mm inner diameter, Sutter Instruments Co., Novato, CA, USA) were pulled into patch pipettes with 2-5 MΩ resistance on a Flaming-Brown Micropipette Puller (Sutter Instruments Co., Novato, CA, USA).

For experiments measuring electrophysiological properties and intrinsic and current-injected excitability, patch pipettes were filled with a potassium-gluconate (K-gluc)-based internal solution containing (in mM) 135 K-gluc, 5 NaCl, 2 MgCl_2_, 10 HEPES, 0.6 EGTA, 4 NA_2_ATP, 0.4 NA_2_GTP, pH 7.35, 290 mOsm. A total of 55 cells were used for analysis (CON M: n = 13, N = 4; EtOH M: n = 16, N = 5; CON F: n = 10, N = 3; EtOH F: n = 16, N = 5).

First, voltage-current relationships were assessed in voltage clamp by measuring the current response to increasing negative 10 mV steps from a holding potential of -70 mV to -120 mV. Membrane resistance was calculated as the average from the smallest two hyperpolarizing sweeps. Capacitance was calculated by dividing the transient charge across the first 30 ms (Q) elicited by the first hyperpolarizing step by the step size (10 mV). The hyperpolarization-activated depolarizing current (I_h_) was measured by subtracting the steady-state current from the peak initial current elicited by the last hyperpolarizing step (-50 mV). Current density was calculated by dividing the change in current for each step by the cell capacitance. Inward rectification was measured by dividing the difference in the change in current elicited by the largest 10 mV step by the change in current elicited by the smallest 10 mV step: [(I_-50 step_ - I_-50 baseline_) – (I_-40 step_ - I_-40 baseline_)] / [(I_-10 step_ - I_-10 baseline_) – (I_0 step_ - I_0 baseline_)]. Resting membrane potential (RMP)/basal state of neuronal activity were evaluated in current-clamp configuration. Current-injected firing of action potentials using current steps and ramps at both RMP and a common potential of -70 mV were then measured in neurons that were resting at these potentials. Voltage sag was assessed in the hyperpolarization step that plateaued at approximately -85 mV, prior to engagement of inward rectifying K channels, calculated as the difference between the largest hyperpolarization voltage within 100 ms of the step minus the plateau voltage at the end of the step. Time to hyperpolarization plateau was assessed using the step that elicited a plateau of approximately -90 mV. Because of a technical issue with the voltage reporting during current clamp recordings from female cohort 6b, we were only able to collect sufficient data to present in males.

For synaptic transmission experiments, pipettes were filled with a cesium-methanesulfonate (Cs-meth)-based solution containing (in mM) 135 Cs-methanesulfonate, 10 KCl, 10 HEPES, 1 MgCl_2_, 0.2 EGTA, 4 Mg-ATP, 0.3 GTP, 20 phosphocreatine, pH 7.35, 290 mOsm. Lidocaine *N*-ethyl bromide (1 mg/ml) was included in the intracellular solution to block postsynaptic sodium currents. Neurons were held at -55 mV to isolate glutamatergic synaptic transmission and measure spontaneous EPSCs (sEPSCs) followed by +10 mV to isolate GABAergic synaptic transmission and record spontaneous IPSCs (sIPSCs) within individual neurons. A total of 55 cells were used for sPSC analysis (CON M: n = 14, N = 6; EtOH M: n = 15, N = 8; CON F: n = 9, N = 3; EtOH F: n = 17, N = 5). Tetrodotoxin (TTX, 500 nM) was included in the bath aCSF to eliminate action potential-dependent spontaneous neural transmission to measure miniature EPSCs (mEPSCs) and IPSCs (mIPSCs). A total of 48 cells were used for mPSC analysis (CON M: n = 15, N = 6; EtOH M: n = 12, N = 7; CON F: n = 8, N = 3; EtOH F: n = 13, N = 4). Electrophysiological recordings of synaptic transmission were used to determine the frequency and amplitude of these synaptic events, which were used to calculate excitatory and inhibitory synaptic drive (PSC frequency x amplitude), as well as the synaptic drive ratio (excitatory synaptic drive / inhibitory synaptic drive). Voltage-current relationships were also assessed in the cells in which sPSCs were measured to confirm that Cs blocked K^+^ and HCN channel-mediated currents assessed using the K-Gluc intracellular recording solution.

Signals were digitized at 10 kHz and filtered at 3 kHz using a Multiclamp 700B amplifier and analyzed using Clampfit 10.7 software (Molecular Devices, Sunnyvale, CA). Traces were excluded from analysis of specific measures if there was spontaneous activity or instability in the section of trace needed for analysis.

### 2.4 Statistical analysis

For all measures, data distributions within group were examined for normality in raw and log space. As we have previously found in monkey and mouse BNST, synaptic transmission measures, membrane resistance and capacitance, and I_h_ were lognormally distributed; for these measures, values are presented in their raw form on a log2 scale y axis in graphs and analyzed following log transformation. All other measures were normally distributed in raw space and so analyzed as such and presented with linear y axes. We identified outliers with Q-Q plots and Rout tests (Q = 1%). Only one outlier value was identified (in inward rectification score) and excluded from analysis; no other values for any measure were excluded. Two-way ANOVAs (2xANOVAs) and two-way and three-way repeated measures ANOVAs (2x and 3xRM-ANOVAs) were used to evaluate the effects of sex and chronic alcohol intake on electrophysiological measures. Significant main effects and interactions were further probed with post hoc direct comparisons using unpaired t-tests corrected for multiple corrections using the Holm-Sidak correction and reported with their adjusted p values. Fisher’s exact tests were used to compare proportions of neurons in states of excitability vs. rest and proportions of neurons that entered depolarization block between groups. One-sample t-tests were used to evaluate whether synaptic drive ratios significantly differed from 1.0. All statistical tests were two-tailed with an α value of 0.05. All data are presented as mean +/- standard error of the mean and individual data points are included where possible in graphs.

## 3. Results

Here, we examined how sex contributes to the basal function of neurons in the rhesus monkey BNST, as well as how long-term voluntary alcohol consumption differentially affects this basal function. Monkeys trained to voluntarily consume EtOH were allowed to consume 4% EtOH for 22 hr/day (termed “open access”) for approximately one year prior to necropsy (**Fig. 1A,B**). A block of brain tissue containing one hemisphere of the BNST and surrounding tissue (**Fig. 1C**) was dissected and sliced into 250 um sections for whole-cell patch-clamp electrophysiological recordings. Using a K-Gluc-based intracellular recording solution, the voltage-current relationship was measured using -10 mV steps from a holding potential of -70 mV to -120 mV (**Fig. 1D**). We found that females had reduced current responses to hyperpolarization of the membrane potential compared to males, and that a history of EtOH drinking decreased these current responses (**Fig. 1E,F**). While the capacitance of BNST neurons did not differ across groups (**Fig. 1G**), current responses normalized to capacitance (current density) remained smaller in females and decreased in EtOH compared to CON groups within sex, especially at hyperpolarized potentials (**Fig. 1H**).

We found that membrane resistance was higher in females than males, and EtOH exposure did not impact this property (**Fig. 2A**). The hyperpolarization-activated depolarizing current (I_h_) measured was lower in female than in male rhesus BNST neurons, and EtOH exposure decreased I_h_ in both sexes (**Fig. 2B**). In addition, male BNST neurons displayed greater inward rectification than those in females, and a history of EtOH decreased inward rectification in both sexes (**Fig. 2C**). We further assessed the relationship between these measures, finding that the I_h_ magnitude and inward rectification score were inversely correlated with membrane resistance (**Fig. 2D and E**, respectively) and positively correlated with one another (**Fig. 2F**). Thus, cells with low I_h_ tended to have high membrane resistance and little inward rectification, suggesting they may be particularly excitable in response to excitatory synaptic inputs. Using a cesium-based intracellular recording solution to block voltage-gated K^+^ channels, including voltage-gated inward rectifier K^+^ (K_ir_) channels, and HCN channels mediating I_h_, we found that current responses to membrane hyperpolarization and I_h_ were minimal and were similar across groups (**Fig. 2G-I**), confirming the involvement of these channels in the effects we observed. These results suggest that female BNST neurons may respond to changes in current across the membrane with greater voltage change, and that they have fewer currents actively maintaining RMP and counteracting depolarizing currents, such as HCN and K_ir_. Further, chronic EtOH drinking may decrease the expression and/or function of these membrane-stabilizing channels, leading to increased excitability.

**Figure 2:**
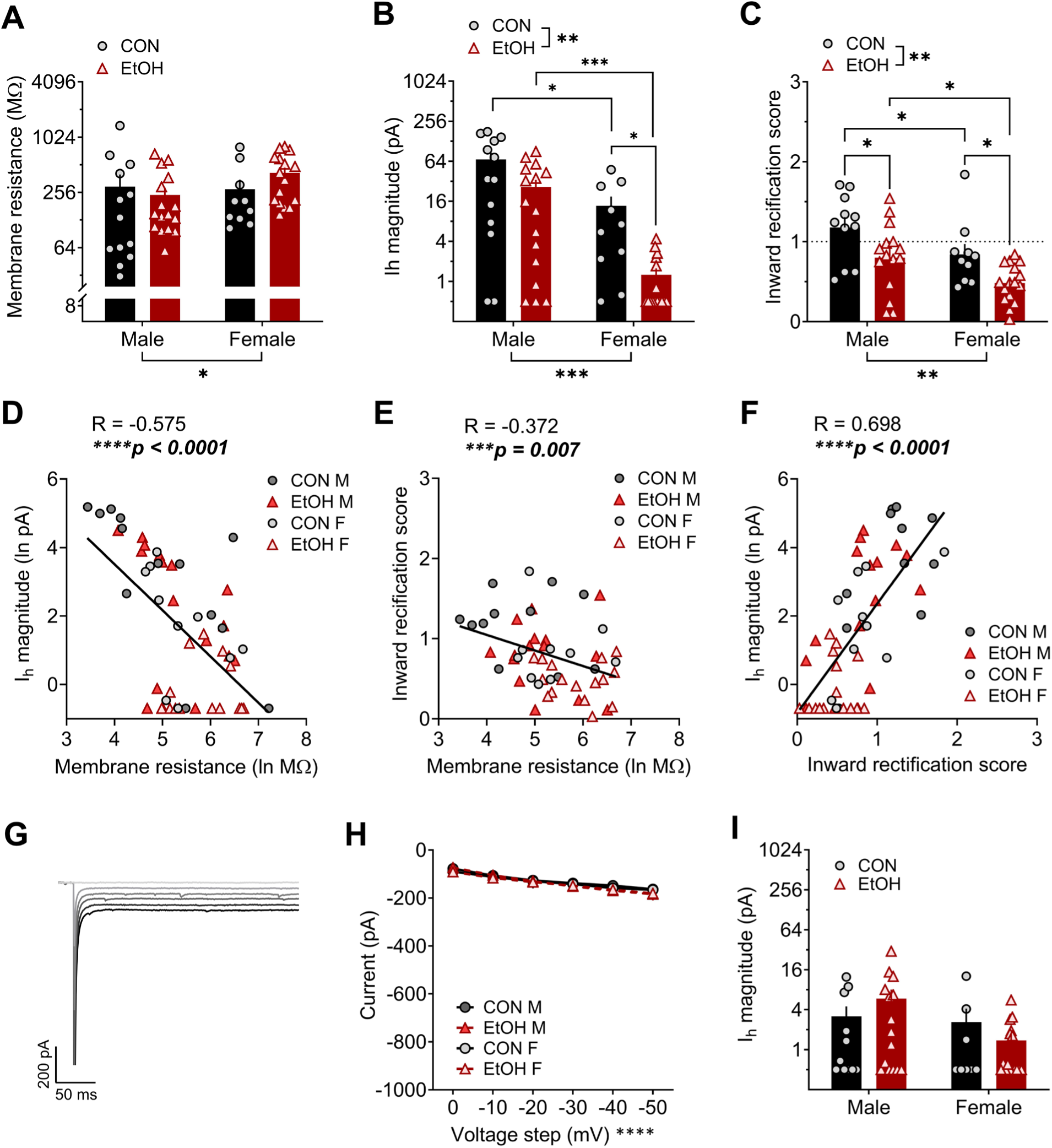
Voltage-gated channel function in monkey BNST neurons differs between males and females and is modulated by a history of chronic voluntary alcohol intake. A) Membrane resistance was higher in female BNST neurons than those of males when measured during voltage clamp recordings. 2xANOVA: main effect of sex (F_1,51_ = 4.84, *p = 0.032) but no effect of EtOH or interaction (ps > 0.10). Post hoc direct comparisons showed a trend for a difference between alcohol males (EtOH M) and females (EtOH F; t_51_ = 2.23, p = 0.060) and no difference between control males (CON M) and females (CON F; p = 0.320). **B)** The hyperpolarization-activated depolarizing current (I_h_) was lower in females than males and decreased by a history of EtOH exposure. 2xANOVA: main effects of sex (F_1,51_ = 16.40, ***p = 0.0002) and EtOH (F_1,51_ = 10.03, **p = 0.003) but no interaction (p > 0.30). Post hoc t-tests showed a significant difference between M and F in both the CON (t_51_ = 2.03, *p = 0.048) and EtOH (t_51_ = 3.88, ***p = 0.001) groups, as well as between CON and EtOH F (t_51_ = 2.80, *p = 0.014) but not M (t_51_ = 1.64, p = 0.108). **C)** Inward rectification was higher in males and decreased by EtOH exposure. 2xANOVA: main effects of sex (F_1,48_ = 9.67, **p = 0.003) and EtOH (F_1,48_ = 12.14, **p = 0.001) but no interaction (p = 0.934). Post hoc direct comparisons showed a significant sex difference between CON M and F (t_48_ = 2.07, *p = 0.044) and EtOH M and F (t_48_ = 2.38, *p = 0.042) and effect of EtOH in M (t_48_ = 2.54, *p = 0.028) and F (t_48_ = 2.39, *p = 0.028). **D-F)** Membrane resistance was negatively correlated with I_h_ in BNST cells (**D**; R = -0.575, ****p < 0.0001) and inward rectification score (**E**; R = -0.372, ***p < 0.007); I_h_ and inward rectification score were highly positively correlated (**F**; R = 0.698, ****p < 0.0001). **G-I)** Effects of sex and EtOH history on voltage-current relationships (as measured in Fig. 1D**,E**) were ablated when K and HCN channels were blocked using a Cs-based intracellular recording solution. Representative trace from a CON M BNST neuron (**G**) and quantification (**H**) of the voltage-current. 3xRM-ANOVA: main effect of voltage step (F_5,230_ = 83.83, ****p < 0.0001) but no other effects or interactions (ps > 0.35). **I)** I_h_ was diminished when a Cs-based intracellular recording solution was used, ablating effects of sex and EtOH. 2xANOVA: no effects or interaction (ps > 0.10).

We used current-clamp recordings to evaluate basal population level excitability, resting membrane potential (RMP), and current-injected firing in these BNST neurons in male monkeys (data in female monkeys not shown due to a technical issue during data collection). We found that a minority of BNST neurons were in a basal state of firing activity in CON M (33%; **Fig. 3A,B**), and a history of EtOH drinking did not affect this basal population-level excitability (**Fig. 3A**) or the RMP of basally inactive/resting neurons (**Fig. 3C**). We further examined whether I_h_ and membrane resistance were related to basal activity state, collapsed across CON M and EtOH M groups because of the low number of basally excited neurons sampled (**Fig. 3D-G**). We found that I_h_ was higher in resting than active neurons and negatively correlated with RMP within resting neurons (**Fig. 3D,E**); membrane resistance was lower in resting than active neurons and positively correlated with RMP within resting neurons (**Fig. 3F,G**). When a ramp current injection protocol was used to elicit action potential firing from RMP in neurons basally at rest (**Fig. 3H**), the membrane potential at which the first action potential was elicited was similar between CON M and EtOH M (**Fig. 3I**), however the minimum current injection needed to elicit that action potential (rheobase) was lower in EtOH M than CON M (**Fig. 3J**). Interestingly, RMP and rheobase were negatively correlated (**Fig. 3K**), showing that while RMP did not significantly differ between CON M and EtOH M monkey BNST neurons, a slightly depolarized RMP in EtOH M neurons may contribute to their greater excitability at RMP. Supporting this, when all neurons were held at a common membrane potential of -70 mV, there was no difference in the rheobase between CON M and EtOH M BNST neurons (**Fig. 3L**).

**Figure 3:**
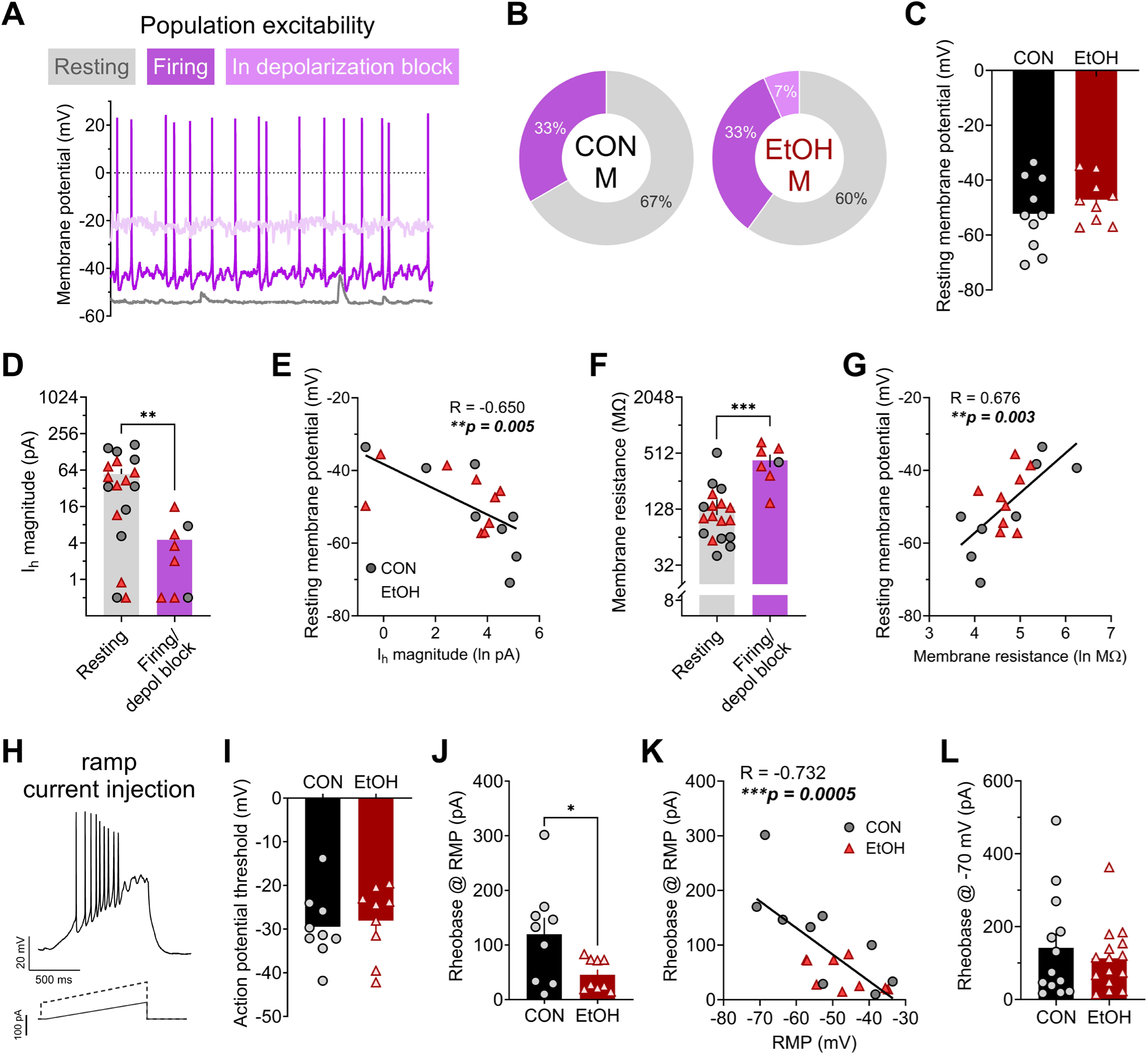
Effects of chronic alcohol drinking on current-injected firing in male rhesus BNST neurons. **A)** Illustration of various basal states of activity observed in male rhesus BNST neurons measured in current clamp configuration. **B)** The proportions of neurons in each basal state did not differ between control males (CON M) and alcohol males (EtOH M; Fisher’s exact test: p > 0.99). **C)** The resting membrane potential (RMP) of neurons basally at rest did not differ between CON M and EtOH M (unpaired t-test: t_17_ = 1.01, p = 0.327). **D-E)** I_h_ magnitude was higher in resting neurons compared to those basally firing or in depolarization block (**D**; unpaired t-test: t_24_ = 3.10, **p = 0.005) and negatively correlated with RMP in resting neurons (**E**; Pearson’s correlation: R = -0.650, **p = 0.005). F-G) Membrane resistance was lower in resting neurons compared to those basally firing or in depolarization block (**G**; unpaired t-test: t_23_ = 4.69, ***p = 0.0001) and negatively correlated with RMP in resting neurons (**E**; Pearson’s correlation: R = -0.650, **p = 0.005). **H-K)** Action potential firing in response to a ramp current injection stimulus in current clamp at RMP. **H)** Example trace for (above) and stimulus waveform of (below) a ramp current-injected firing during current-clamp recordings in BNST neurons at RMP. **I)** The threshold to fire an action potential did not differ between CON M and EtOH M (unpaired t-test: t_16_ = 0.38, p = 0.708). **J)** The rheobase (minimum amount of injected current required to elicit an action potential) measured from RMP in resting neurons was lower in EtOH M than CON M (unpaired t-test with Welch’s correction: t_9.6_ = 2.34, *p = 0.042). **K)** Rheobase was negatively correlated with RMP in male BNST neurons (R = -0.732, ***p = 0.0005). **L)** When neurons were held at a common membrane potential of -70 mV, there was no difference in rheobase between CON M and EtOH M (unpaired t-test: t_26_ = 0.64, p = 0.527).

We also examined action potential firing from -70 mV using a step current injection protocol (**Fig. 4A,B**), finding that BNST neurons in CON M and EtOH M fired a similar number of action potentials in response to increasing depolarizing current steps (**Fig. 4C**) and a similar, small proportion of neurons entered a state of depolarization block, a putative homeostatic mechanism to prevent excitotoxicity, during this protocol (**Fig. 4D**). Within this protocol, we then evaluated the presence of a T-type calcium current (I_t_) in neurons upon depolarization, as indicated by prolonged depolarization following action potentials and/or doublets (illustrated in **Fig. 4E**), which regulates membrane potential in response to initial depolarization near RMP. We found that a similar proportion of BNST neurons in CON M and EtOH M monkeys displayed I_t_, (**Fig. 4F**) and that I_t_+ neurons were similarly likely to be basally active as I_t_- neurons (**Fig. 4G**). We also examined whether hyperpolarizing current injection steps could elicit a transient voltage sag (**Fig. 4A,H**), which has been used as a proxy measure for I_h_ in current clamp recordings and has been used to categorize BNST cell types along with measures of inward rectification and I_t_ (Hammack et al., 2007). Similar to a previous publication in monkeys, we found that rhesus male BNST neurons displayed little to no voltage sag (Daniel et al., 2017) regardless of alcohol history, with all cells displaying values below 1 mV (where the presence of sag has been defined as above 2 mV; **Fig. 4H,I**).

**Figure 4:**
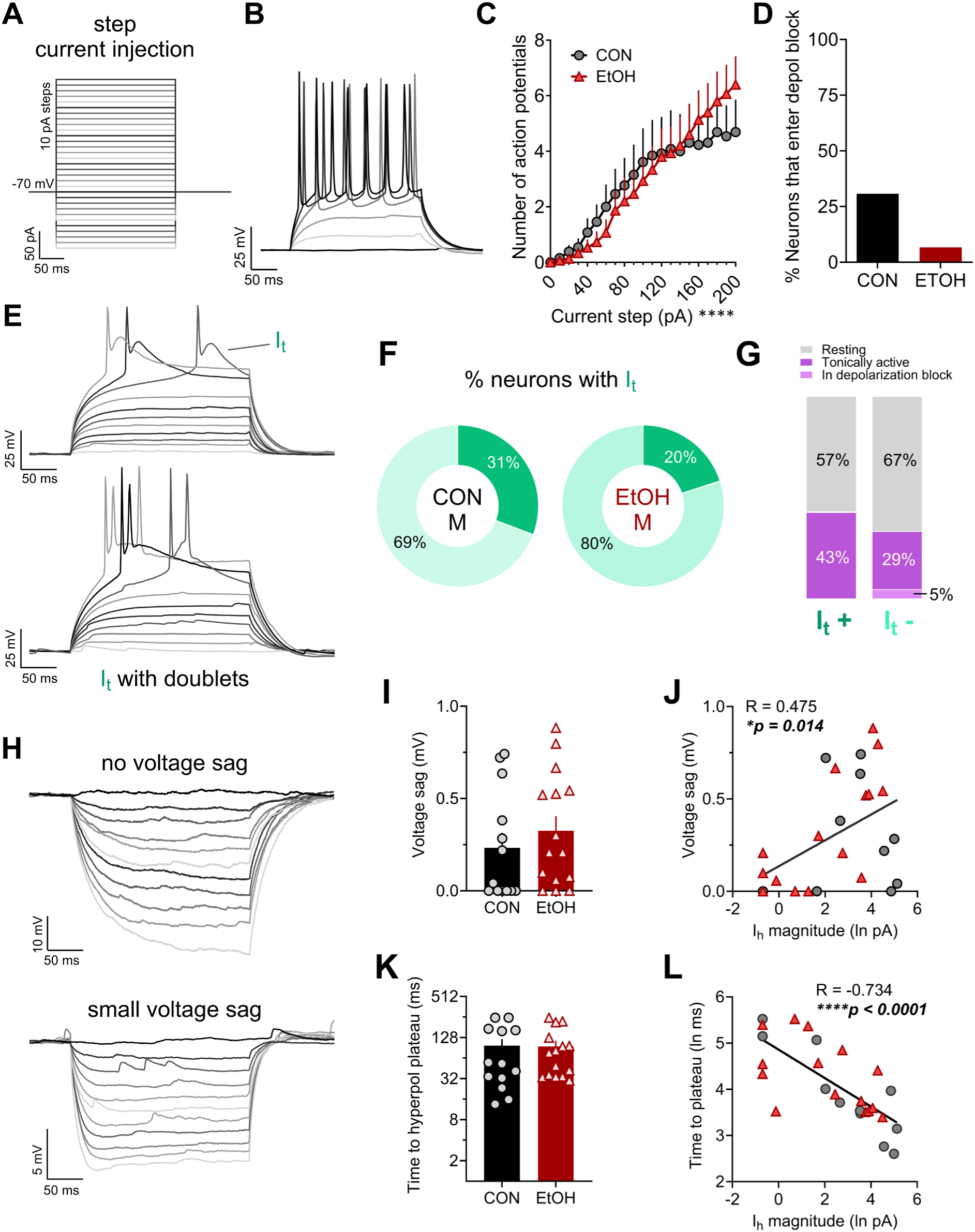
Voltage responses of male rhesus BNST neurons to step current injection during current clamp recordings. A) Current injection protocol used at a holding potential of -70 mV, in which 10 pA steps from -100 to +200 were used across sweeps to elicit hyper- and depolarizing responses. **B)** Representative trace of responses to increasingly depolarizing current steps (every 4^th^ step from 0 to 20 pA shown). **C)** EtOH did not affect the number of action potentials fired in response to step current injection. 2xRM-ANOVA: main effect of current step (F_20,520_ = 25.79, ****p < 0.0001) but no effect of EtOH or interaction (ps > 0.30). **D)** The proportion of BNST neurons that entered depolarization block during step current injection was not different between CON M and EtOH M (Fisher’s exact test: p = 0.153). **E)** Representative traces illustrating a voltage-gated calcium T-type current (I_t_)-mediated prolonged depolarization following a single action potential (top) and eliciting a doublet (bottom) during depolarizing steps (every other step shown). **F)** The proportion of neurons displaying I_t_ was similar in CON M and EtOH M BNST neurons (Fisher’s exact test: p > 0.670). **G)** The proportion of I_t_+ neurons that were basally active was similar in CON M and EtOH M BNST neurons (Fisher’s exact test: p > 0.674). **H**) Representative traces of responses to hyperpolarizing current steps, showing one cell that displayed no voltage sag and a long time to reach the hyperpolarization plateau (top) and another cell that displayed a small voltage sag at the beginning of the hyperpolarization steps (bottom). **I-J)** The magnitude of voltage sag was small and did not differ in BNST neurons from CON M and EtOH M (**I**: unpaired t-test: t_26_ = 0.81, p = 0.423), however it was positively correlated with I_h_ measured in voltage clamp (**J**: Pearson’s correlation: R = 0.475, *p – 0.014). **K-L)** The time to hyperpolarization plateau at the step reaching approximately -90 mV did not differ between CON M and EtOH M (**K**; unpaired t-test: t_26_ = 0.47, p = 0.644) but was negatively correlated with I_h_ magnitude (**L**; Pearson’s correlation: R = -0.734, ****p < 0.0001).

However, voltage sag magnitude was weakly correlated with I_h_ magnitude measured in voltage clamp (**Fig. 4J**). Because of the relative absence of sag, we further assessed the time to reach hyperpolarization plateau at the step that elicited a potential of approximately -90 mV (prior to eliciting inwardly rectifying K^+^ channels), in which there was variability across cells. While alcohol history did not affect this measure (**Fig. 4K**), it was highly negatively correlated with I_h_ magnitude within cells (**Fig. 4L**). Thus, time to hyperpolarization plateau was a better current clamp correlate of I_h_ magnitude than voltage sag, suggesting that other significant conductances regulating membrane potential are at play in rhesus BNST neurons that may compete with those mediated by HCN channels upon hyperpolarization, an idea suggested previously for mouse BNST neurons (Miura et al., 2022). Altogether, these excitability data provide converging evidence that increased excitability in response to depolarization in EtOH M compared to CON M was driven primarily by voltage-gated channel function regulating resting membrane potential.

Next, we evaluated whether synaptic transmission in rhesus BNST neurons were affected by sex or altered by alcohol drinking. First, we evaluated direct, local spontaneous synaptic transmission by measuring mPSCs when network-dependent activity was prevented by the sodium channel blocker TTX (500 nM) in the extracellular bath aCSF (**Fig. 5**, with example traces in **Fig. 5A**) and found that mEPSC, but not mIPSC, measures were affected. mEPSC frequency was higher in females than males and EtOH slightly increased this measure (trend for effect; **Fig. 5B**), while mIPSC frequency was similar across all groups (**Fig. 5C**). However, neither mEPSC nor mIPSC amplitude were affected by sex or EtOH (**Fig. 5D and E**). Together, this led to higher mPSC excitatory synaptic drive in females than males and a trend for increased excitatory drive by EtOH (**Fig. 5F**). In contrast, there were no differences in mIPSC synaptic drive (**Fig. 5G**). Therefore, the composite measure of excitatory/inhibitory synaptic drive ratio (SDR) was higher in females than males, indicating a higher bias toward excitation in females; again, there was a trend for an effect of EtOH in this measure, leading overall to a ratio significantly below 1.0 in CON M and significantly above 1.0 in EtOH F (**Fig. 5H**).

**Figure 5:**
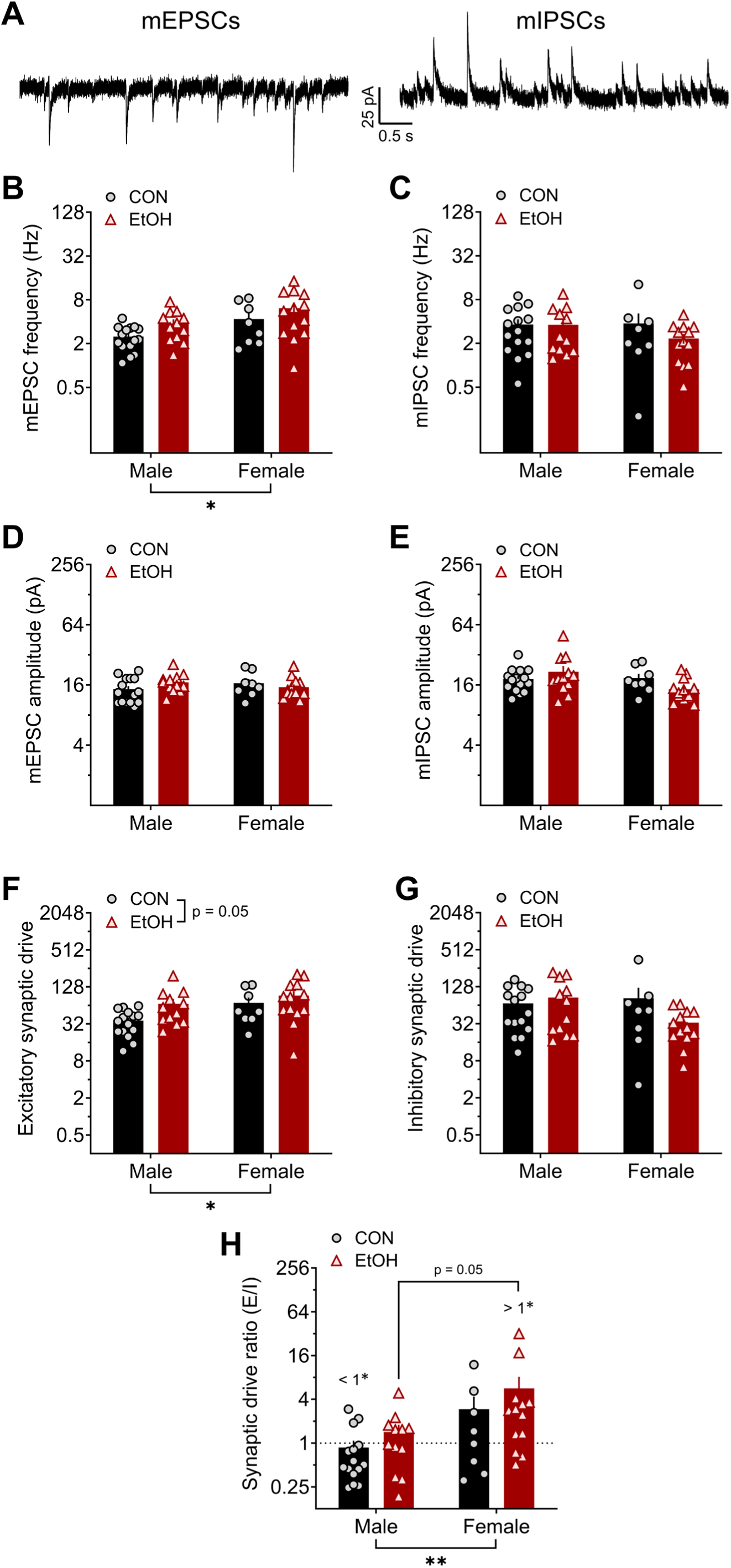
Excitatory, but not inhibitory, local synaptic transmission in BNST neurons is affected by sex and a history of alcohol drinking: miniature postsynaptic currents (mPSCs). A) Representative traces of excitatory mPSCs (mEPSCs; left) and inhibitory mPSCs (mIPSCs; right) from an individual BNST neuron measured at holding potentials of -55 mV and 10 mV, respectively, from control males (CON M) and females (CON F) and alcohol males (EtOH M) and females (EtOH F). **B)** mEPSC frequency was higher in females than males (2xANOVA: main effect of sex (F_1,44_ = 5.36, *p = 0.025), and there was a trend of EtOH (F_1,44_ = 3.87, p = 0.056) but no interaction (p > 0.75). Post hoc t-tests revealed no significant differences between males and females in either CON or EtOH monkeys (ps > 0.15). **C)** There were no differences in mIPSC frequency (ps > 0.20). **D-E)** Neither mEPSC (**D**) nor mIPSC (**E**) amplitude were affected by sex or EtOH (ps > 0.05). **F)** Excitatory synaptic drive (mEPSC frequency x amplitude) was higher in females and increased by chronic alcohol exposure. 2xANOVA: main effects of sex (F_1,44_ = 4.62, *p = 0.037) and EtOH (F_1,44_ = 4.02, p = 0.051) but no interaction (p > 0.35). Posthoc t-tests revealed no significant differences in direct comparisons (ps > 0.05). **G)** Inhibitory synaptic drive was unaffected by sex and EtOH. 2x ANOVA: no effects (ps > 0.10). **H)** The synaptic drive ratio (excitatory synaptic drive / inhibitory synaptic drive) was higher in females than males. 2xANOVA: main effect of sex (F_1,44_ = 8.13, **p = 0.007), trend for effect of EtOH (F_1,44_ = 3.07, p = 0.087), and no interaction (p > 0.75). Posthoc t-tests showed a trend for higher SDR in EtOH F than EtOH M (t_44_ = 2.31, p = 0.050) and in CON F than CON M (t_44_ = 1.75, p = 0.087). in addition, CON M had a ratio below 1.0 (one-sample t-test: t_14_ = 2.26, ^#^p = 0.041), indicating a net synaptic inhibition, and EtOH F had a ratio above 1.0 (one-sample t-test: t_12_ = 2.87, ^#^p = 0.014), while CON F and EtOH M ratios did not differ from 1.0 (ps > 0.45).

When network-dependent synaptic transmission was left intact during recordings of spontaneous PSCs (sPSCS), inhibition was primarily affected by sex and EtOH (**Fig. 6**). Further, although sEPSC frequency was unaffected by sex or EtOH (**Fig. 6A**), sIPSC frequency was higher in BNST neurons from EtOH compared to CON monkeys (**Fig. 6B**). This finding is consistent with our published work in male rhesus monkeys and male mice with a history of binge alcohol drinking that suggested an upregulation of GABAergic interneuron activity in the BNST following chronic alcohol exposure (Pleil et al., 2015a; Pleil et al., 2015b; Pleil et al., 2016). The amplitude of both sEPSCs (**Fig. 6C**) and sIPSCs (**Fig. 6D**) was lower in BNST neurons from females than males, especially in EtOH monkeys. sPSC excitatory synaptic drive was not altered by sex or EtOH (**Fig. 6E**), but sPSC inhibitory synaptic drive was higher in males than females and increased by EtOH (**Fig. 6F**). This led to an excitatory/inhibitory synaptic drive ratio below 1.0, indicating a net synaptic inhibition of BNST neurons, in EtOH males but no other groups (**Fig. 6G**).

**Figure 6:**
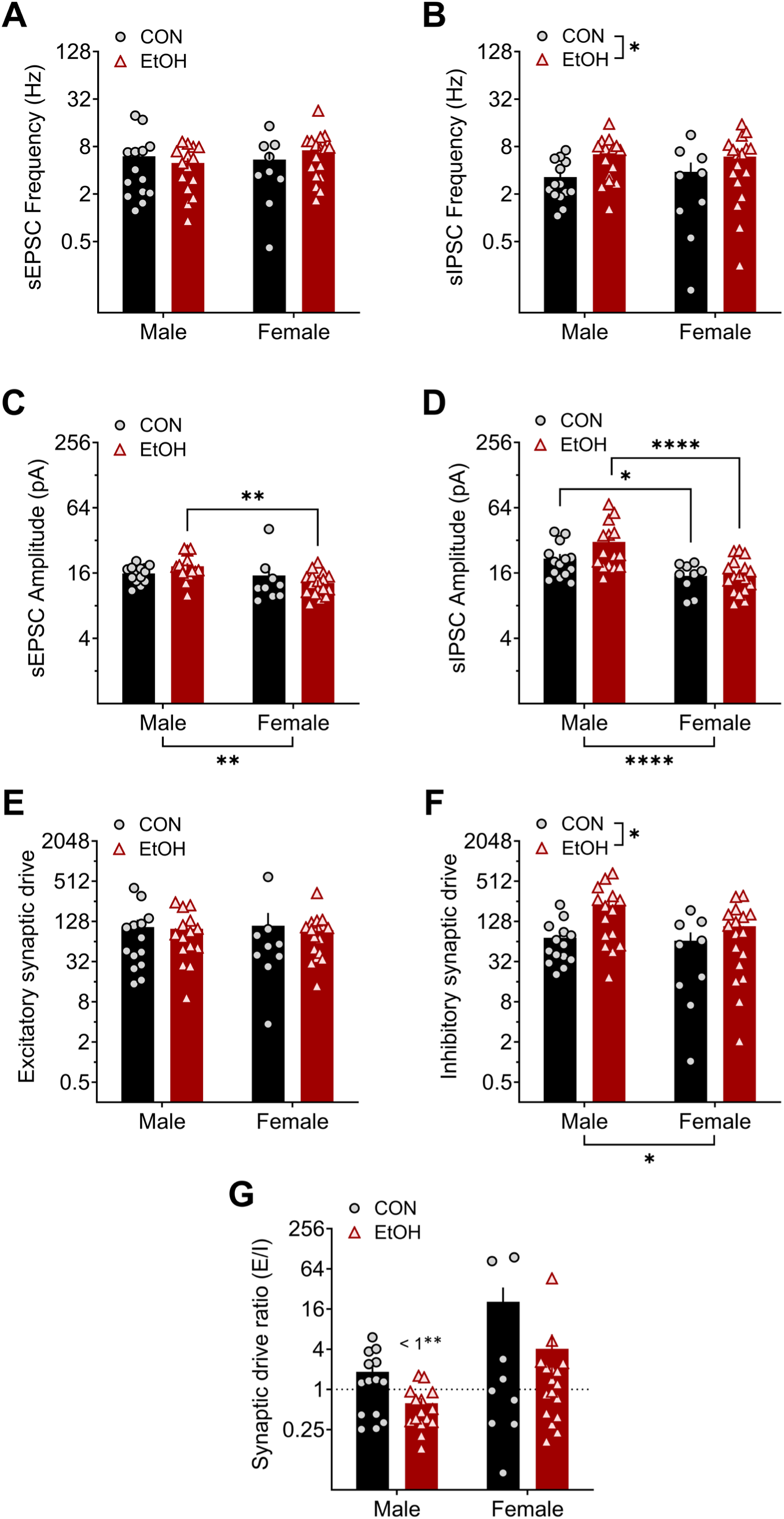
A history of alcohol drinking and sex modulate activity-dependent synaptic transmission in BNST neurons: spontaneous postsynaptic currents (sPSCs). A) Spontaneous excitatory postsynaptic current (sEPSC) frequency was unaltered by sex and alcohol. 2xANOVA: no effects (ps > 0.30). **B)** Spontaneous inhibitory postsynaptic current (sIPSC) frequency was increased by alcohol exposure. 2xANOVA: main effect of EtOH (F_1,51_ = 6.47, *p = 0.014) but no effect of sex or interaction (ps > 0.25). Posthoc t-tests between CON and EtOH groups within sex: ps > 0.10. **C)** sEPSC amplitude was lower in females than males, an effect drive by the EtOH group. 2xANOVA: main effect of sex (sex: F_1,51_ = 9.95, **p = 0.003) but no effect of EtOH or interaction (ps > 0.20). Post hoc t-tests between M and F within group: CON (t_51_ = 1.22, p = 0.228), EtOH (t_51_ = 3.47, **p = 0.002). **D)** sIPSC amplitude was lower in females than males in both CON and EtOH conditions. 2xANOVA: main effect of sex (sex: F_1,51_ = 20.28, ****p < 0.0001) but no effect of EtOH or interaction (ps > 0.10). Post hoc t-tests between M and F within group: CON (t_51_ = 2.09, *p = 0.042), EtOH (t_51_ = 4.53, ****p < 0.0001). **E)** Spontaneous excitatory synaptic drive was not affected by sex or alcohol exposure. 2xANOVA: no effects (ps > 0.35). **F)** Spontaneous inhibitory synaptic drive was higher in males than females and increased by a history of alcohol exposure. 2xANOVA: main effect of sex (F_1,51_ = 5.25, *p = 0.026) and EtOH (F_1,51_ = 6.15, *p = 0.017) but no interaction (p > 0.70). Post hoc t-tests revealed no significant differences (ps > 0.05). **G)** The spontaneous synaptic drive ratio was not different between groups. 2xANOVA: no effects (ps > 0.10). However, EtOH M had a ratio below 1.0 (one-sample t-test: t_14_ = 3.85, ^##^p = 0.002), indicating a net synaptic inhibition, while other groups did not differ from 1.0 (ps > 0.50).

## 4. Discussion

Here, we found that female rhesus BNST neurons have higher membrane resistance as well as less inward rectification and I_h_, currents mediated predominantly by voltage-gated K+ and HCN channels, respectively, than those in males. These results suggest that female BNST neurons may be more excitable than male BNST neurons. A history of alcohol exposure also decreased these currents in both sexes and slightly increased the excitability of male BNST neurons (females not assessed). We also found that males had higher synaptic inhibition when network activity was left intact, and alcohol drinking increased inhibition in both sexes. This led to a synaptic drive ratio biased toward inhibition in BNST neurons of male alcohol-drinking monkeys. In contrast, excitatory synaptic drive was higher in females when network activity was blocked, and alcohol exposure tended to increase this synaptic excitation as well.

### 4.1 Homeostatic inhibition as a key feature of the BNST response to chronic alcohol

We found that chronic alcohol exposure increased spontaneous IPSC frequency, a putative measure of GABA release, onto BNST neurons in both male and female rhesus BNST neurons. These results are consistent with our previous observation of this effect in male rhesus monkeys and in mice (Pleil et al., 2015b; Pleil et al., 2016). As these previous studies did not include female subjects, our observation of a similar effect on synaptic inhibition in female primate BNST fills an important gap in the literature. Further, here we assessed miniature PSCs and found that excitation, but not inhibition was affected by alcohol exposure, suggesting that increased glutamate release from excitatory efferents into the BNST may drive a network activity-dependent increase in synaptic inhibition by the local interneuron population. We previously reported in mice that the synaptic input from the paraventricular thalamus, the densest source of glutamatergic input to the BNST that robustly engages the inhibitory microcircuit in the BNST, is upregulated following a history of alcohol (Levine et al., 2021). It may be the case that the PVT (or other excitatory inputs to the BNST) and the inhibitory neurons that they synapse onto are particularly sensitive to alcohol-induced plasticity. Future studies are needed to assess the role of excitatory inputs to the BNST in alcohol-related behaviors and the circuit organization underlying alcohol-induced plasticity in both rodents and monkeys. Further, the variability in sIPSC frequency in female BNST neurons compared to those in males that we observed here points to potential sex differences in underlying interneuron microcircuit organization of the BNST that require additional investigation of sex differences in primate BNST micro- and macro-circuit organization.

We also found that in both sexes, alcohol exposure decreased the function of currents important for regulating resting membrane potential and limiting action potential generation in response to depolarization of the membrane, including inward rectification mediated primarily by voltage-gated inward rectifier K^+^ (K_ir_) and I_h_ by HCN channels. Therefore, the chronic alcohol-induced increase in network activity-dependent synaptic inhibition may be a homeostatic response to increased excitability in BNST neurons to limit their activity. In males, we found that a history of alcohol drinking contributed to increased sensitivity to current injected-firing (as measured by rheobase). While RMP was not different between control and alcohol-exposed monkey neurons, RMP and rheobase were highly correlated. Further, membrane resistance was highly correlated with I_h_, RMP, and rheobase, suggesting that effects of alcohol on HCN channels may contribute to differences in the membrane potential and sensitivity to excitation (via current injection) measured at RMP observed between CON M and EtOH M. Indeed, when cells were held at a common membrane potential, this sensitivity to current did not differ between groups. This suggests that a reduction in the currents controlling membrane potential contributed to increased excitability in males; it is further possible that increased excitatory synaptic input participates in the inhibition of HCN and other voltage-sensitive channels. There is a great deal of evidence from rodent studies that calcium-gated and voltage-gated ion channels regulating membrane potential and neuronal excitability, including HCN, Kv7, K_ir_, and small-conductance Ca^2+^-activated K^+^ (SK) channels, promote alcohol drinking and are susceptible to alcohol-induced plasticity (Welsh et al., 2011; Padula et al., 2015; Rinker et al., 2017; Salling et al., 2018). Additionally, a recent study in rhesus monkeys that included subjects in our study showed that SK channels that regulate neuronal firing are dysregulated in the nucleus accumbens, an adjacent and interconnected extended amygdala structure, in heavy alcohol drinking monkeys (Mulholland et al., 2023). Future studies are needed to directly assess the roles of these and other ion channels in the effects of alcohol exposure on excitability in rhesus BNST neurons and to determine the relationship between synaptic and excitability effects of sex and alcohol.

### 4.2 Variation in electrophysiological features in BNST neurons within and across species

BNST neurons in rodents have previously been categorized into multiple types based on voltage responses to hyperpolarizing and depolarizing current injection steps, including inward rectification and voltage sag, the latter of which has been used as a proxy measure for HCN channel-mediated I_h_, and action potential firing pattern (Type I: large voltage sag, little inward rectification, persistent firing, and no I_t_; Type II: little/no voltage sag or inward rectification, large I_t_, burst firing, and/or rebound firing; Type III: large inward rectification, little/no voltage sag or I_t_) (Hammack et al., 2007; Daniel et al., 2017). While these categories were determined in male rats and segregate BNST neurons into equal thirds in that subject population, both male mouse and male monkey BNST neurons have been shown to be less distinguishable based on the combination of these measures (Silberman et al., 2013; Daniel et al., 2017; Miura et al., 2022). This is, in part, due to the presence of additional features in these cells, such as irregular firing and a lack of voltage sag in most neurons (which we also found here). Further, in male mice, the use of these categories has not proven to be useful in distinguishing specific cell types of BNST projection populations. For example, both the CRF neuron and VTA-projecting neuron populations in the BNST of male mice have been shown to have all three Types, as well as a large proportion of neurons that fall into an “Other” category because they do not fit into the three Types (Silberman et al., 2013; Miura et al., 2022). Here, the use of voltage clamp recordings (in the same cells in which we performed current clamp recordings) to more directly assess I_h_ showed than many cells in CON M monkeys displayed a large I_h_ in voltage clamps even though they displayed almost no voltage sag in current clamp. Thus, while there was a large effect of EtOH on decreasing I_h_, this effect was not detected when using voltage sag as a proxy measure. Similarly, a comprehensive study in male mice that also measured I_h_ in voltage clamp and voltage sag in current clamp showed that these measures were not related to one another, finding that VTA-projecting BNST neurons displayed any possible combination of these (large I_h_ and large sag, large I_h_ but no sag, no/little I_h_ but large sag, and no I_h_ or sag) (Miura et al., 2022). Altogether, these data suggest that voltage sag is not a suitable proxy measure for I_h_ in monkey (and perhaps mouse) BNST neurons, likely because there are other conductances activated at hyperpolarized potentials that interfere with this as a measure or contribute to its presence, as well as the ability to detect effects, such as alcohol-mediated plasticity in our study. And, while we were able to detect putative Type II neurons that displayed I_t_, we found that I_h_ and inward rectification score (both measured in voltage clamp) were positively correlated in monkey BNST cells, while Types I and III have classically been defined by their mutually exclusive expression based on current clamp measures. Therefore, regardless of whether these differences are driven by the inconsistency in the relationship between voltage and current clamp measures or fundamental species differences in the display of voltage sag, the diversity of electrophysiological properties in BNST neurons suggest that alternative features may need to be considered when trying to classify these in male monkeys and mice. Future investigation is required to determine how these categories may be useful for distinguishing female BNST neurons in any species.

### 4.3 Independent effects of sex and alcohol on BNST neuron function

We observed several interesting sex differences in the basal function of BNST neurons in rhesus monkeys. For example, we found that female BNST neurons displayed higher membrane resistance, less inward rectification, and almost no I_h_ compared to male neurons that displayed robust rectification and I_h_. While we were unable to assess excitability in current clamp recordings in females, these differences suggest BNST neurons may be more basally excitable in naïve females than males. We also saw that direct synaptic excitation of BNST neurons was greater in females than males; both of these observations are similar to those we have made in the mouse BNST (Levine et al., 2021). While we found here that putative glutamate release and alcohol-induced increase in direct synaptic excitation were higher in females, network activity-dependent synaptic inhibition, primarily due to polysynaptic inhibition engaged by glutamatergic inputs, was not higher in females. This suggests that the female BNST network does not engage a homeostatic inhibitory network to counter the higher excitability and synaptic excitation. As BNST neuron excitability and synaptic excitation have been shown to contribute to increased alcohol drinking, anxiety-like and depressive-like behavior in rodents (Silberman et al., 2013; Pleil et al., 2015b; Levine et al., 2021), underlying sex differences in the excitability of BNST neurons may confer increased susceptibility to the expression of these behaviors even in the absence of alcohol exposure. Given the observation in primate brain here, these may translate to humans and be related neuropsychiatric disease susceptibility in females (Grant et al., 2004a; Grant et al., 2004b; Kessler et al., 2005; Peltier et al., 2019; Flores-Bonilla and Richardson, 2020; Guinle and Sinha, 2020).

While we found many sex differences in BNST neuron function and independent effects of alcohol exposure on these neurons, we found few interactions between these factors. That is, alcohol often affected male and female BNST neurons similarly even when there were basal sex differences in function. In particular, we saw that the effects of alcohol were similar in both sexes in the currents we measured that affect intrinsic excitability and glutamate release; interestingly, this occurred in a manner that shifted the cellular phenotype in the direction that females were basally compared to males. That is, alcohol increased direct synaptic excitation, which was basally higher in females, and reduced membrane potential-stabilizing K^+^ and HCN currents, which were basally lower in females. These results highlight the possibility that female BNST neurons have a phenotype that is basally more susceptible to the hyperexcitability effects of alcohol on BNST function, perhaps as lower membrane-stabilizing I_h_ promotes the postsynaptic excitability in response to excitatory input. In contrast, chronic alcohol exposure also increased network activity-dependent synaptic inhibition in both sexes that was already basally higher in males than females. This resulted in a meaningful overall shift toward synaptic inhibition of BNST neurons following alcohol exposure in males but not females, a phenomenon thought to be an adaptive homeostatic mechanism to counter alcohol-induced hyperexcitability. These results are similar to those we have found in mice, in which the excitability of a subpopulation of stress-sensitive BNST neurons and glutamate release (from PVT excitatory inputs) onto them were increased following a history of alcohol exposure in males to the basally higher levels observed in females, and males met this hyperexcitability with increased inhibition (Levine et al., 2021). As high BNST activity promotes alcohol drinking, anxiety, and negative affect and is associated with related neuropsychiatric diseases, these underlying sex differences in BNST function that bias females more toward hyperexcitability and males toward synaptic inhibition may confer greater susceptibility in females to alcohol-induced plasticity that contributes to maladaptive behavioral states.

Furthermore, the measures of synaptic function we assessed here tended to be more variable in female than male rhesus BNST neurons, suggesting there may also be sex differences in the circuit organization underlying this sex-dependent circuit function and differential neuromodulation of these BNST circuits in rhesus monkeys. We recently showed that signaling of ovarian-derived estrogen in the BNST drives synaptic excitation and alcohol drinking in female mice (Zallar et al., 2023). This mechanism may undergo plasticity as a consequence of chronic alcohol drinking that contributes to the sex-dependent effects we found here. Other studies have examined the effects of long-term alcohol consumption on the circulating levels of sex and stress hormones in this nonhuman primate model, and we and others have examined the relationship between these hormones and the function of the BNST and other brain regions in males (Pleil et al., 2016; Dozier et al., 2019). Future consideration of hormone dysregulation in females may be key to better understanding their reciprocal relationships with alcohol drinking in translational models of alcohol drinking.

## Author contributions

KEP designed slice electrophysiology experiments, collected and analyzed the data, and wrote the manuscript. TLK designed and oversaw slice electrophysiology experiments. VCCC oversaw slice electrophysiology experiments. KAG designed and oversaw all *in vivo* alcohol drinking manipulations and data collection. All authors edited and approved the manuscript.

